# Caveolin-1 controls lineage fidelity of adult neural stem cells via lysosomal degradation of PDGFRα

**DOI:** 10.64898/2025.12.16.694780

**Authors:** He Zhang, Yusuke Kihara, Mitsuho Sasaki, Takuya Tomita, Yasushi Saeki, Taeko Kobayashi

## Abstract

Caveolae are cell-surface signaling hubs involved in stem cell proliferation and differentiation. However, the molecular details of how they influence stem cell quiescence remain poorly understood. In this study, we show that most caveolae components, particularly caveolin-1 (Cav-1), are specifically upregulated in quiescent neural stem cells (NSCs). Furthermore, we demonstrate that Cav-1 restricts platelet-derived growth factor receptor (PDGFR) α signaling, a key regulator for NSC proliferation and oligodendrocyte differentiation in the adult brain, through lysosomal degradation. Genetic ablation of Cav-1 leads to aberrant NSC activation, reduced neurogenesis, and enhanced ectopic oligodendrocyte differentiation on the ventricular surface in the ventral side of the ventricular-subventricular zone (V-SVZ), a key NSC niche responsible for neuroblast differentiation. Mechanistically, we show that Cav-1 deficiency suppresses the endo-lysosomal degradation of PDGFRα, leading to PDGFRα accumulation and excessive signaling for oligodendrocyte differentiation. Taken together, our results uncover a crucial homeostatic mechanism wherein caveolae function as a safeguard to maintain adult NSC quiescence and lineage fidelity by controlling lysosomal turnover of PDGFRα in a niche-specific manner.

## Introduction

In the adult mammalian brain, neurogenesis continuously occurs throughout life in two specific regions: the ventricular-subventricular zone (V-SVZ) of the lateral ventricles and the subgranular zone (SGZ) of the dentate gyrus (DG) in the hippocampus (Altman & Das, 1965; Doetsch et al, 1999; Eriksson et al, 1998). These neurogenic areas have specialized microenvironments known as stem cell niches, where neural stem cells (NSCs) can regulate their self-renewal and fate decisions, differentiating into neurons, astrocytes, and oligodendrocytes (Kjell et al, 2020; Lim et al, 2000; Silva-Vargas et al, 2016; Urban et al, 2019). In these niches, most adult NSCs remain quiescent, a reversible arrest at the G0/G1 phase of the cell cycle that maintains the long-term regenerative capacity of the stem cell pool and prevents premature exhaustion, ensuring a continuous supply of new neurons during adulthood (Kalamakis et al, 2019; Urban et al, 2019). Recent studies have shown that the regional differences in these niches reflect the heterogeneity of NSCs and their distinct fate choices (Bond et al, 2015; Cebrian-Silla et al, 2021). In the V-SVZ of the lateral ventricles, the dorsal (top) and ventral (bottom) positions of the V-SVZ differentially influence their neurogenic or oligodendrogenic fates: ventral NSCs exhibit a strong bias toward neuroblast-generating fates, while dorsal NSCs exhibit a propensity towards oligodendrogenic fates (Cebrian-Silla et al, 2021; Ortega et al, 2013; Radecki & Samanta, 2022; Willis et al, 2025). However, the regulatory mechanism behind this location-specific fate determinations has not been fully elucidated.

Caveolae, which are cholesterol- and sphingolipid-rich, flask-shaped invaginations on the cell membrane, are essential microdomains for signal transduction. They serve multiple functions, acting as plasma membrane organizers, stress protectors, and signal regulators that integrate mechanical and chemical cues (Badaut et al, 2024; Couet et al, 1997; Engelman et al, 1998; Liu et al, 2002; Parton, 2018; Yamamoto et al, 1999). Caveolae participate in lipid rafts, organize and cluster signaling molecules, mediate clathrin-independent endocytosis for their lysosomal degradation, and contribute to cholesterol homeostasis. Caveolin-1 (Cav-1), an integral membrane protein, is essential for caveolae formation (Drab et al, 2001). It oligomerizes with Cavin, the caveolae-associated protein, to create the characteristic caveosome structure (Hill et al, 2008). Cav-1 binds to and regulates various signaling factors, holding them in inhibitory complexes and releasing them upon stimulation (Pelkmans et al, 2004). Biochemical studies have shown that the N-terminal scaffolding domain of Cav-1 directly interacts with short motifs in target proteins, controlling their activity (Engelman et al, 1998).

Cav-1 has been reported to be involved in an altered balance between self-renewal and differentiation in tissue stem cells (Baker & Tuan, 2013). Cav-1 knockout (KO) results in increased activation of quiescent hematopoietic stem cells, leading to the transient expansion followed by exhaustion of the stem cell pool (Bai et al, 2014). In mesenchymal stem cells, Cav-1 influences lineage commitment, shifting differentiation from adipogenic toward osteogenic fate upon Cav-1 depletion (Bandara et al, 2016). In the adult brain, Cav-1–KO mice exhibit accelerated neurodegeneration and aging-related phenotypes, suggesting defects in adult NSC maintenance (Tang et al, 2021; Trushina et al, 2006). Some studies suggest that Cav-1 differentially affects NSC behavior depending on their two NSC niches, the DG and the V-SVZ. In the DG, Cav-1–KO increases the number of newly generated neurons with high VEGF expression levels in 9-week–old mice (Li et al, 2011c), enhances oligodendrocyte differentiation associated with low β-catenin signaling (Li et al, 2011a), and reduces the number of astroglial cells with low Notch signaling (Li et al, 2011b). Cav-1 conditional knockout (cKO) using Nestin-Cre-ERT2 mice results in decreased hippocampal NSC proliferation cell-autonomously in 3- and 6-month-old mice and increased neurogenesis in the DG in 6-month-old mice, along with changes in mitochondrial fusion in the NSCs (Stephen et al, 2023). On the other hand, in the V-SVZ, Cav-1–KO increases the number of proliferating NSCs in 8-week–old mice (Jasmin et al, 2009). These findings highlight the multiple functions of Cav-1 involving in several signaling pathways for adult NSC behavior and guiding appropriate proliferation and differentiation of adult NSCs, and that the functional roles of Cav-1 highly depends on the two niches. However, the molecular mechanisms underlying these niche-dependent roles of Cav-1 remain poorly understood.

We recently reported that quiescent NSCs upregulate lysosomal factors, which actively maintain their quiescent state (Kobayashi et al, 2019), and that lysosomal proteolytic activity fluctuates with age in adult NSCs (Zhang et al, 2024). These findings raise the possibility that a potential delivery system involved in lysosomal proteolysis may also contribute to NSC maintenance in the adult brain. Here, we investigated NSC quiescence regulators using proteomics approaches and identified that caveolae components are more abundant in quiescent NSCs compared to proliferating NSCs. Our findings further suggest that caveolae play NSC niche- and age-dependent roles in the adult mouse brain, differentially affecting NSC proliferation and fate determination in the V-SVZ. We further showed that Cav-1 negatively regulates platelet-derived growth factor receptor alpha (PDGFRα), a key receptor on oligodendrocyte precursor cells (OPCs), via lysosomal degradation. These results propose that caveolae function as a crucial regulator of NSC quiescence and lineage fidelity of NSCs in the adult mouse brain.

## Results and Discussion

### Cav-1 upregulation during neural stem cell transition to quiescence

To identify potential regulators of NSC quiescence, we performed a tandem mass tag (TMT)-based proteomic analysis of proliferating (active NSCs: aNSCs) and quiescent (qNSCs) NSCs *in vitro* (Morita et al, 2025). Quiescence was induced by switching from proliferating to quiescent medium and culturing for at least 3 days (Martynoga et al, 2013). Proteome data indicated a global change in protein components between aNSCs and qNSCs (Fig. 1A, Table EV1A). We searched for components specifically enriched in qNSCs (Fig. 1B, Table EV1B). Gene ontology analysis using qNSC-high proteins showed that many plasma-membrane-related terms including caveola were significantly enriched in qNSCs (Fig. 1C, Table EV2). Cav-1, Caveolin-2 (Cav-2), and caveolae-associated proteins Cavin1 and Cavin3, as well as caveolae formation-related proteins Pacsin1, Pacsin2 and Pacsin3, were significantly upregulated in qNSCs compared to aNSCs (Fig. 1A,B,D) (Hansen et al, 2011; Parton et al, 2021), suggesting that caveolae play a role on regulating NSC quiescence. In these caveolae-related factors, Cav-1, Cavin3, and Pacsin3 showed increased mRNA levels as well as protein levels in qNSCs compared to aNSCs (Fig. 1E, Table EV3). Cav-1, Cavin3, and Pacsin3 were upregulated 7.6-, 8.5-, and 2.3-fold at the protein level, respectively, and 18-, 1.9-, and 2.4-fold at the mRNA level, respectively, in qNSCs compared to those in aNSCs. These results demonstrate that qNSCs increase the expression of these caveolae-related factors at both the mRNA and protein levels. Increased Cav-1 expression during transition from proliferating to quiescence was also observed by both immunocytochemistry (Fig. 1F, day 1-3) and western blotting (Fig. 1G,H, day 1-3). In immunocytochemistry, Ki-67–negative, non-proliferative NSCs showed high levels of Cav-1 (Fig. 1F, day1-2), suggesting that Cav-1 may be involved in the regulation of quiescence in adult NSCs.

**Figure 1.**
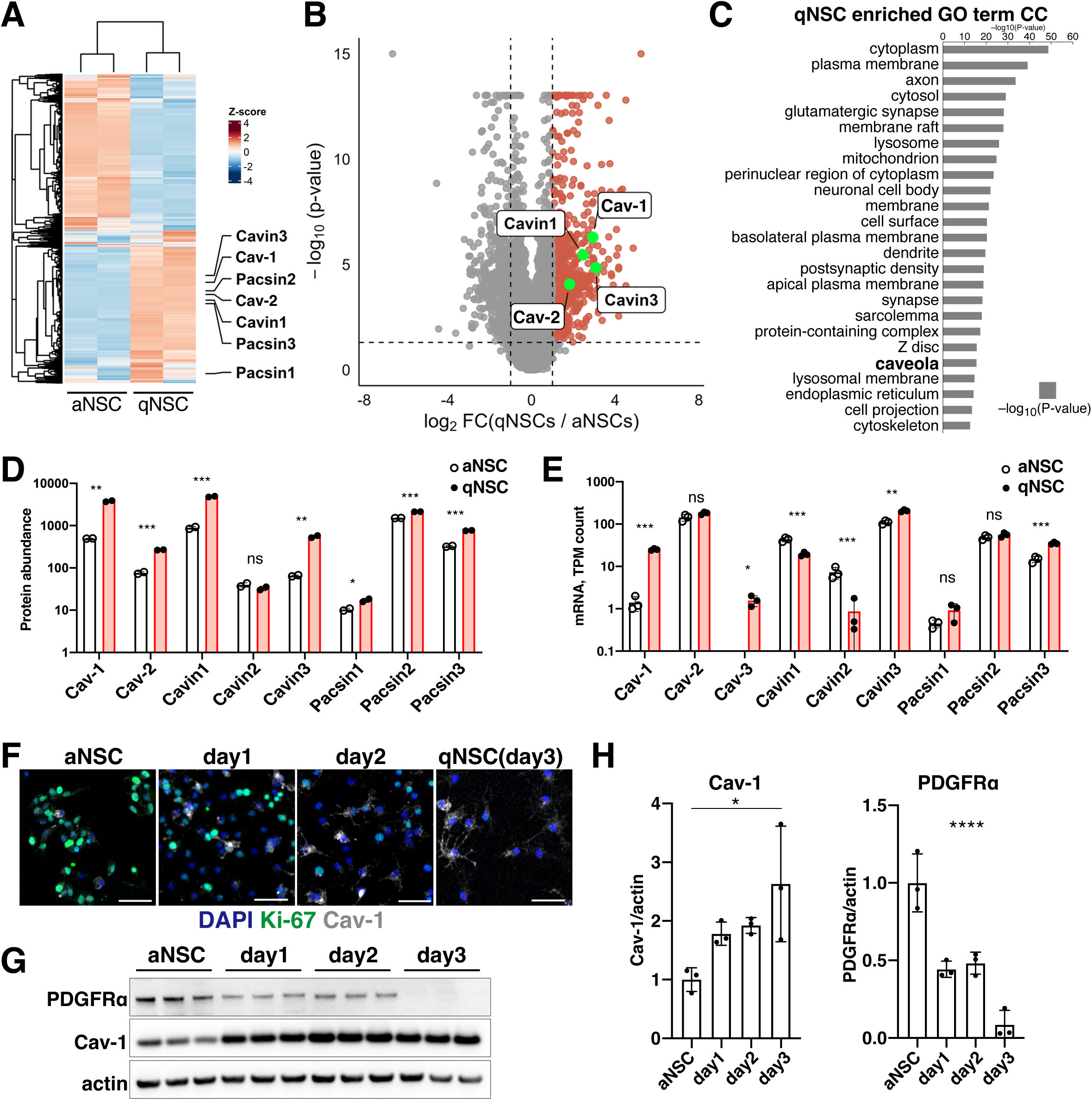
Caveolae-related proteins are enriched in quiescent NSCs. (A) Quantitative TMT–mass spectrometry of proliferating NSCs (aNSCs) and quiescent NSCs (qNSCs) reveals overall proteome changes caused by quiescence. Heatmap shows z-score–normalized protein abundance across samples. (B) Volcano plot of qNSC vs. aNSC proteomics. Vertical dashed lines indicate |log₂FC| = 1; horizontal dashed line marks p = 0.05. (C) Gene ontology cellular component enrichment of proteins upregulated in qNSCs. (D) Quantification of caveolae-related protein abundance in aNSCs and qNSCs using TMT-MS. Data are plotted on a log-10 scale and presented as mean ± SD; n = 3 per group and independent preparations; unpaired two-tailed t-test (ns, not significant; *p < 0.05; **p < 0.01 and ***p < 0.001). Cav-3 was not detectable in our TMT-MS. (E) Quantification of caveolae-related mRNA abundance from RNA-seq. TPM values are plotted on a log-10 scale and presented as the mean ± SD; n = 3 per group; DESeq2, (ns, not significant; *Padj < 0.05; **Padj < 0.01 and ***Padj < 0.001). (F) Immunocytochemistry showing Cav-1 (white) and Ki-67 (green) expression during the transition from aNSCs to qNSCs. Nuclei are labeled with DAPI (blue). Scale bars, 100 µm. (G) Western blots of Cav-1 and PDGFRα during quiescence induction. Actin was used as a loading control. (H) Quantification of Cav-1/actin and PDGFRα/actin ratios from (G). Average of Cav-1/actin and PDGFRα/actin in aNSC was set to 1 in the bar chart, respectively. Data are shown as the mean ± SD; n = 3 per group; one-way ANOVA (PDGFRα) with Tukey’s multiple-comparison test (Cav-1) (*p < 0.05 and ****p < 0.0001).

### Cav-1–KO mice exhibit niche-dependent phenotypic differences in the SVZ

We investigated the role of Cav-1 in adult NSCs of the mouse brain by analyzing two neurogenic niches in Cav-1–KO mice, which have a single base insertion in exon 3 introduced via clustered regularly interspaced short palindromic repeats (CRISPR)–CRISPR-associated protein 9 (Cas9), at various ages: two-week–old (P14), two-month–old (2MO), and six-month–old (6MO). In the adult brains of heterozygous (HET) control mice, Cav-1 protein appeared as spots in both the cell bodies and processes of NSCs and astrocytes in the DG and V-SVZ (Fig. EV1A,B), and its expression declined with age in NSC niches, with a significant age-related decrease observed in NSCs of the V-SVZ and the DG astrocytes (Fig. EV1C). These results are consistent with the previously reported age-related reduction in Cav-1 expression in the hippocampus (Head et al, 2010). To evaluate the role of Cav-1 in adult NSC quiescence and neurogenesis in the brain, we examined NSC number and proliferation and newly generated neurons in the DG and V-SVZ of Cav-1–KO mice (Fig. EV2 and Fig. 2). In the DG, Cav-1 deletion did not affect any NSC behavior compared to Cav-1–HET mice in P14 or 6MO mice (Fig. EV2A,B,E,F). Cav-1–KO 2MO mice exhibited decreased numbers of proliferating (aNSCs: Sox2+Nestin+Ki-67+) and total (NSCs: Sox2+Nestin+) NSCs (Fig. EV2C,D), but the proportion of proliferating NSCs in total NSCs (aNSCs%) remained unchanged (Fig. EV2B,D,F). Newly generated neurons (DCX+) in the DG of Cav-1–KO mice increased in 6MO mice (Fig. EV2I,J), but did not change in 2MO mice (Fig. EV2G,H), compared to those in Cav-1–HET mice. These results suggest that the loss of Cav-1 does not affect NSC activation, but does affect neurogenesis at 6MO in the DG.

**Figure 2.**
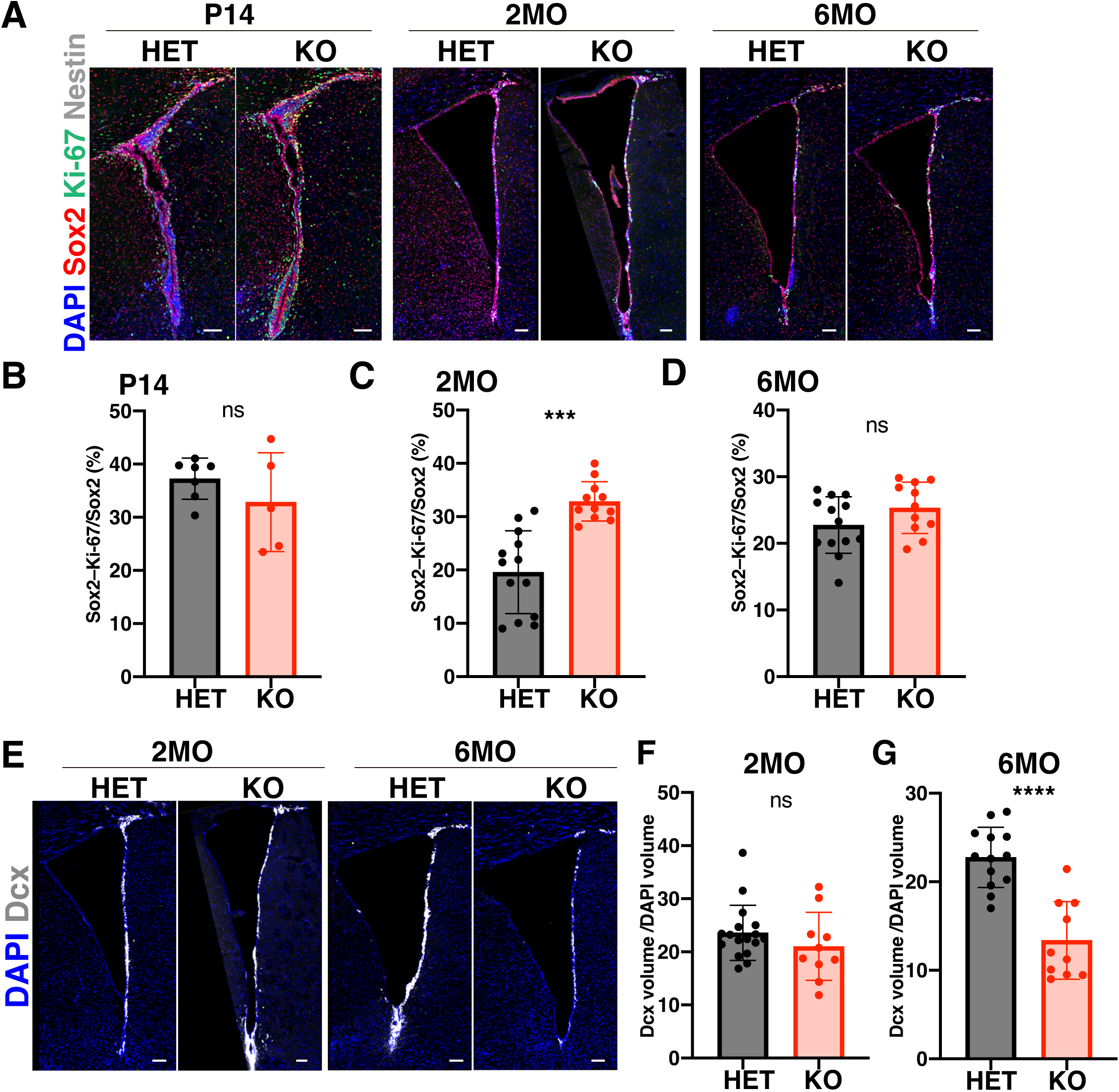
Cav-1 deletion increases NSC activation but reduces neurogenesis in the SVZ in an age-dependent manner. (A) Representative confocal images of the V-SVZ in Cav-1–HET and Cav-1–KO mice at postnatal day 14 (P14), 2 months (2MO), and 6 months (6MO). Sox2 (red), Ki-67 (green), and Nestin (white) label NSCs (Sox2⁺) and aNSCs (Sox2⁺Ki-67⁺). Nuclei are labeled with DAPI (blue). Scale bars, 100 µm. (B–D) Quantification of aNSCs (Sox2+Ki-67+) normalized to total NSCs (Sox2+) in the V-SVZ across ages and plotted as percentages. Mean ± SD; n = 3 mice per group; unpaired two-tailed t-test (ns, not significant; ***p < 0.001). (E) Representative images of doublecortin-positive (DCX⁺, white) neuroblast with DAPI (blue) in the V-SVZ at 2MO and 6MO. Scale bars, 100 µm. (F, G) Quantification of neurogenesis in the V-SVZ, shown as total DCX⁺ volume normalized to DAPI⁺ volume. Mean ± SD; n = 3 mice per group; unpaired two-tailed t-test (ns, not significant; ****p < 0.0001).

In the V-SVZ, Cav-1–KO mice showed a marked increase in the aNSC/NSC ratio (aNSCs%; aNSCs: Sox2+Ki-67+, NSCs: Sox2+) compared to Cav-1–HET mice at 2MO (Fig. 2A,C), although no significant difference was observed at P14 and 6MO (Fig. 2A,B,D). The aNSC/NSC ratio (aNSCs%) at 2MO was comparable to that observed at P14 in Cav-1–KO mice (Fig. 2B,C). It indicates that Cav-1–KO mice sustain a higher proportion of proliferating NSCs at 2MO than Cav-1–HET mice (Fig. 2C). On the other hand, the number of newly born neurons was significantly decreased in Cav-1–KO mice compared to those in Cav-1–HET mice at 6MO (Fig. 2E,G), although no significant difference was observed at 2MO (Fig. 2E,F). These results suggest that Cav-1–deficient NSCs are more activated in the V-SVZ and can enter the cell cycle more readily than control NSCs in young adult mice (2MO) (Fig. 2C). However, the increased NSC activation did not lead to neuronal output in the V-SVZ and instead resulted in decreased neural differentiation at later stages (6MO) (Fig. 2G). It suggests that Cav-1–KO NSCs might prefer alternative differentiation pathways in the V-SVZ. Collectively, these findings suggest that Cav-1 plays niche- and age-dependent roles for NSCs, and that its deletion more significantly affects NSC behavior in the V-SVZ.

### Cav-1 deletion promotes ectopic oligodendrocyte precursor differentiation in the V-SVZ

To determine whether Cav-1 contributes to differentiation fates, we isolated NSCs from Cav-1–KO and control HET mouse embryos (Fig. EV3), and examined differentiation into three lineages—neurons, astrocytes, and oligodendrocytes—*in vitro* (Fig. 3). Culturing such NSCs in neuronal differentiation medium induced Tuj-1–positive neurons from both Cav-1–KO and Cav-1–HET control NSCs, with lower differentiation efficiency in Cav-1–KO NSCs than in control NSCs (Fig. 3A,B). Astrocyte differentiation produced highly GFAP-positive astrocytes from both Cav-1–KO and Cav-1–HET NSCs, with higher efficiency in Cav-1–KO NSCs than in controls (Fig. 3C,D). Compared with the controls, Cav-1–KO NSCs showed enhanced oligodendrocyte differentiation and produced more oligodendrocyte precursors (OPCs: Olig2^+^PDGFRα^+^) (Fig. 3E,F). These results indicate that Cav-1–KO NSCs remain multipotent but tend to differentiate into glial cells, astrocytes and OPCs, rather than into neurons.

**Figure 3.**
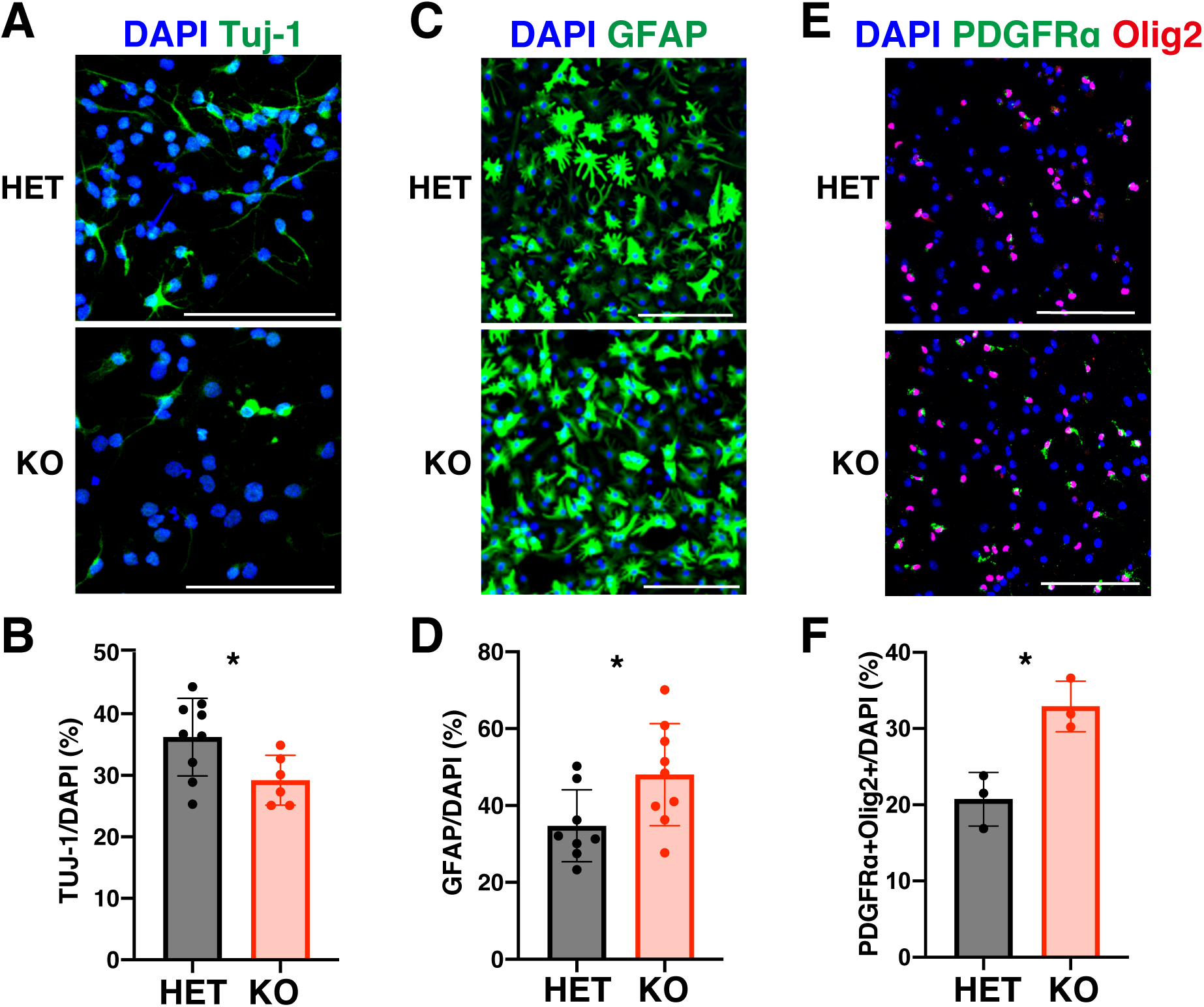
Cav-1 deletion influences NSC differentiation by promoting glial fates and increasing the production of astrocytes and OPCs *in vitro*. (A, C, E) Representative images showing differentiation of Cav-1–HET and Cav-1–KO NSCs into neurons (A), astrocytes (C), and OPCs (E). Neurons were identified by Tuj1⁺ staining (green in A), astrocytes by GFAP⁺ staining (green in C), and OPCs by co-expression of PDGFRα (green in E) and Olig2 (red in E). Nuclei were stained with DAPI (nuclei, blue). Scale bars, 100 µm. (B, D, F) Quantitative analyses of neurons (B; Tuj-1/DAPI), astrocytes (D; GFAP/DAPI), and OPCs (F; PDGFRα⁺Olig2⁺/DAPI) normalized to total DAPI+ cells and shown as percentages. Mean ± SD; n = 3 per group; unpaired two-tailed t-test (*p < 0.05).

The ventral V-SVZ is the area of oligodendrocyte progenitor production from qNSCs *in vivo* (Bond et al, 2015; Cebrian-Silla et al, 2021). We examined OPCs in the V-SVZ of Cav-1–KO mice at 2MO and 6MO of age (Fig. 4A-C). Although the number of Olig2+ cells was unchanged in the lateral V-SVZ at both ages, that of OPCs (Olig2+PDGFRα+) significantly increased at 6MO but not at 2MO in Cav-1–KO mice (Fig. 4B,C). This result provides a complementary mechanism, via enhanced OPC differentiation, for the reduced neurogenesis observed in Cav-1–KO mice at 6MO in the V-SVZ (Fig. 2G). Notably, the number of Olig2+PDGFRα+ OPCs on the ventricular surface of the ventral V-SVZ, known as intraventricular OPCs (Delgado et al, 2021), significantly increased in Cav-1–KO mice at 2MO compared to the control Cav-1–HET mice and remained elevated at 6MO (Fig. 4D-F). To investigate the contribution to astrocyte differentiation, we also examined astrocytes as GFAP-S100β double-positive cells in the V-SVZ of Cav-1–KO and Cav-1–HET mice at P14, 2MO, and 6MO (Fig. EV4A-D). The number of astrocytes remained unchanged between Cav-1–HET and Cav-1–KO mice in the lateral V-SVZ at all ages (Fig. EV4B-D). These results suggest that Cav-1–KO enhances OPC differentiation from adult NSCs in the ventral V-SVZ and induces the production of ectopic intraventricular OPCs.

**Figure 4.**
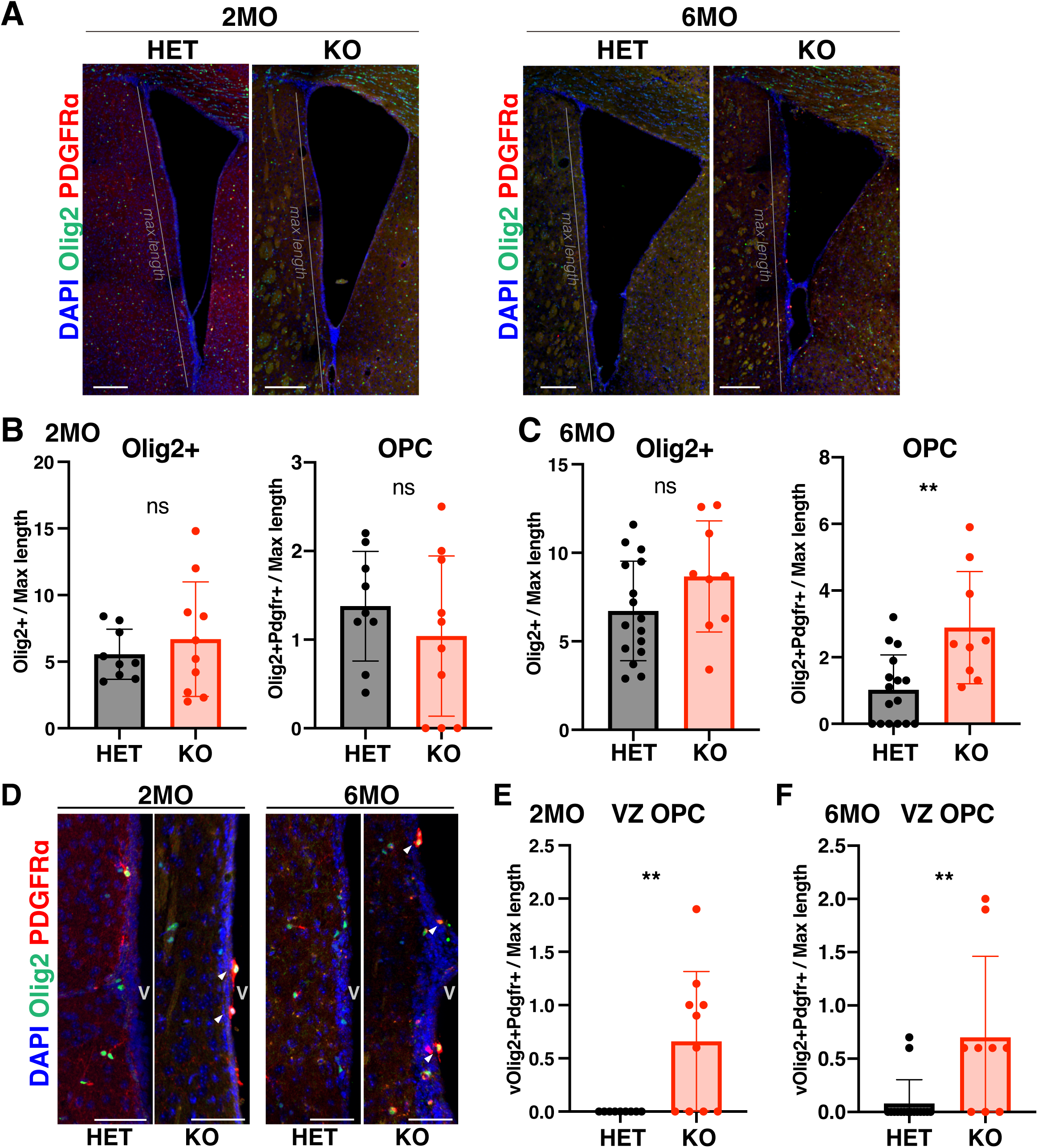
Cav-1 deletion increases OPC production in the V-SVZ and causes ectopic intraventricular OPCs. (A) Representative confocal images of the V-SVZ from Cav-1–HET and Cav-1–KO mice at 2 months (2MO) and 6 months (6MO), stained for Olig2 (green) and PDGFRα (red) to identify OPCs. Nuclei are labeled with DAPI (blue). Grey lines indicate the maximal length of lateral SVZ used for quantification. Scale bars, 200 µm. (B, C) Quantification of oligodendrocyte lineage cells in the V-SVZ at 2MO (B) and 6MO (C), based on the number of Olig2⁺ cells (left) and OPCs (Olig2⁺PDGFRα⁺; right), normalized to the maximal length of the SVZ (A). Mean ± SD; n = 3 mice per group; unpaired two-tailed t-test (ns: not significant; **p < 0.01). (D) High-magnification images of the lateral ventricular side of the V-SVZ at 2MO and 6MO showing ectopic OPCs on the ventricular surface. White arrowheads indicate Olig2⁺PDGFRα⁺ intraventricular OPCs. V denotes ventricle. Scale bars, 50 µm. (E, F) Quantification of intraventricular OPCs (Olig2⁺PDGFRα⁺ cells on the ventricle surface), normalized to the maxima length of the SVZ (A). Mean ± SD; n = 3 mice per group; unpaired two-tailed t-test (**p < 0.01).

### Cav-1 regulates PDGFRα stability via lysosomal degradation in NSCs

To understand the molecular mechanisms underlying NSC behavior in Cav-1–KO mice, RNA sequencing (RNA-seq) analysis of Cav-1–KO and Cav-1–HET NSCs was performed (Fig. 5A, Fig. EV5, Table EV4). We observed significant upregulation of oligodendrocyte-related factors, including PDGFRα and myelin regulatory factor (Myrf) (Fig. 5B) (Kuhn et al, 2019; Bohrer et al, 2015) in Cav-1–KO NSCs, within factors related to NSC differentiation including Wnt4, and S100β (Fig. 5A,B; Fig. EV5A,B). We focused on PDGFRα because the PDGFRα-mediated signaling pathway is known to be crucial for oligodendrocyte differentiation and glioma cell proliferation in the brain (Funa & Sasahara, 2014; Liu et al, 2011). PDGFRα upregulation due to Cav-1 deletion aligns with our *in vitro* and *in vivo* findings of enhanced OPC differentiation (Figs. 3,4) and increased NSC proliferation (Fig. 2) in the V-SVZ. Our western blot results showed that NSC quiescence induction *in vitro* decreased PDGFRα expression while increasing Cav-1 expression (Fig. 1G,H). Previous studies have indicated that some type-B NSCs in the SVZ express PDGFRα and that excessive PDGFRα activation can abnormally increase cell proliferation and ultimately lead to glioma formation (Jackson et al, 2006; Kesari & Stiles, 2006). PDGFRα protein expression is reported to be nearly absent in the DG of the adult brain. In nestin-GFP transgenic mice, which label NSCs, clear colocalization of PDGFRα and GFP-positive cells is detected in the SVZ, whereas such colocalization is not observed in the DG (Gilley et al, 2011). These studies are consistent with our observation that Cav-1–KO exerts milder effects on NSCs in the DG than on those in the V-SVZ. Furthermore, in Cav-1–KO NSCs, PDGFRα protein expression was increased compared to control-HET NSCs (Fig. 5C). The increase in PDGFRα in Cav-1–KO NSCs was 2.1 times at the mRNA (Fig. 5B) and 3.6 times at the protein (Fig. 5C). It implies that the deletion of Cav-1 may affect the post-translational regulation of PDGFRα protein, potentially leading to an increase of PDGFRα mRNA expression via feedback mechanisms of PDGFRα signaling. We investigated whether Cav-1 controls PDGFRα protein levels and stability in NSCs using a cycloheximide chase assay. Cycloheximide is a protein synthesis inhibitor, and the chase assay following the addition of cycloheximide is a widely used method capable of measuring intracellular protein stability kinetics and protein degradation (Li et al, 2021; Zhang et al, 2024). We found that PDGFRα protein degradation kinetics were significantly slowed in Cav-1–KO NSCs, with the half-life of PDGFRα increasing from 1.01 h in control-HET NSCs to 1.84 h in Cav-1–KO NSCs (Fig. 5D). The lysosomal inhibitor bafilomycin A1 suppressed PDGFRα degradation in both control and Cav-1–KO NSCs; however, the proteasomal inhibitor bortezomib did not inhibit PDGFRα degradation in either NSC type (Fig. 5D). These results indicate that Cav-1 negatively regulates PDGFRα protein levels by mediating its lysosomal degradation. Although Cav-1 is known to inhibit PDGFRα function through direct interaction with the caveolin-scaffolding domain of Cav-1 (Yamamoto et al, 1999), our findings reveal an additional layer of regulation in which Cav-1 controls PDGFRα signaling via lysosomal downregulation.

**Figure 5.**
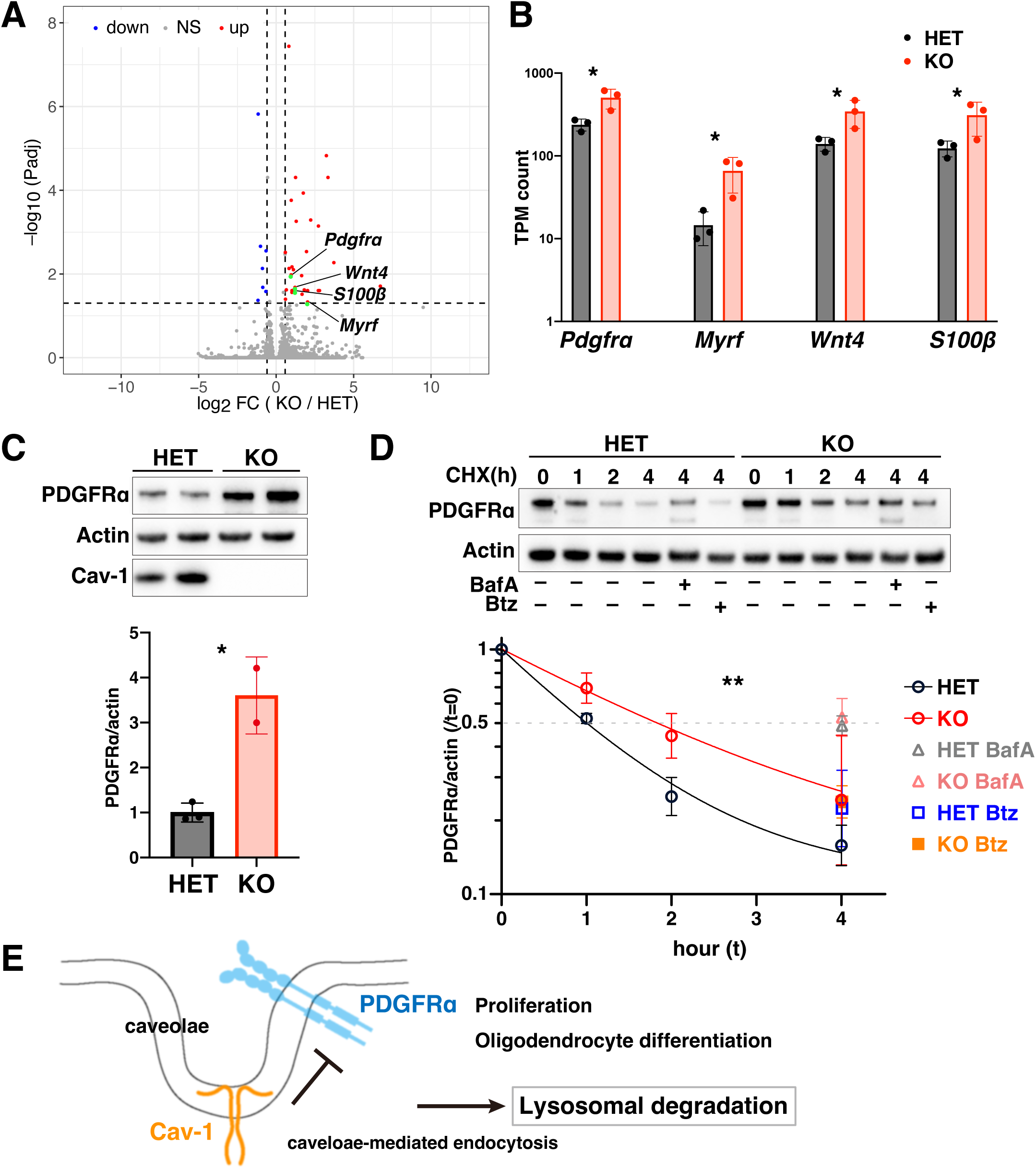
Cav-1 deletion increases PDGFRα expression and slows its lysosomal degradation. (A) Volcano plot of RNA-seq comparing Cav-1–HET and Cav-1–KO NSCs. Vertical and horizontal dashed lines denote fold change = |1.5| and Padj = 0.05, respectively. Blue, red, and grey dots represent genes that are down-regulated, up-regulated, and not significant, respectively. Green dots highlight oligodendrocyte-related genes. (B) mRNA expression levels of Pdgfrα, Myrf, Wnt4, and S100β measured as TPM are plotted on a log-10 scale. Mean ± SD; n = 3 per group; DESeq2 (*Padj < 0.05). (C) Western blot analysis of PDGFRα protein levels in Cav-1–HET and Cav-1–KO NSCs. Actin was used as a loading control. Average band intensity of PDGFRα/actin in HET was set to 1 in the bar chart. Mean ± SD; n = 2 (KO) and 3 (HET); unpaired two-tailed t-test (*p < 0.05). (D) Cycloheximide (CHX) chase assay evaluating PDGFRα degradation at 0, 1, 2 and 4 h in Cav-1–HET (black circles) and Cav-1–KO NSCs (red circles), with (4 h) or without (0, 1, 2 and 4 h) bafilomycin A1 (BafA; gray [HET] and pink [KO] triangles in lower chart) or bortezomib (Btz; blue [HET] and orange [KO] squares in lower chart). Quantified values of PDGFRα normalized to actin are plotted after dividing by the value at t=0 (lower chart). Non-linear fitting used to calculate PDGFRα half-lives in Cav-1–HET (black line) and Cav-1–KO NSCs (red line), plotted with a log-10 scale on the y-axis. Half-lives of PDGFRα are 1.01 h in Cav-1–HET and 1.84 h in Cav-1–KO NSCs (gray dotted line, T1/2). Mean ± SD; n = 3 per group; paired t-test between HET (black circles) and KO (red circles) (**p < 0.01). (E) Proposed model: Cav-1 in caveolae facilitates endo-lysosomal degradation of PDGFRα via clathrin-independent endocytosis, thereby restricting PDGFRα signaling. Loss of Cav-1 results in decreased PDGFRα turnover, increased PDGFRα signaling, enhanced NSC proliferation, and higher ectopic OPC differentiation.

Overall, in this study, we found that Cav-1 levels increase during NSC quiescence and that Cav-1 regulates NSC proliferation and oligodendrocyte precursor differentiation in the V-SVZ. The molecular mechanism underlying these phenotypes is that Cav-1 inhibits PDGFRα signaling by promoting lysosomal degradation of PDGFRα. These results suggest that the caveolae-mediated endo-lysosomal pathway is essential for maintaining and guiding the appropriate differentiation and proliferation of adult NSCs, depending on their niches. Beyond NSC regulation, caveolae are also involved in the regulations of lipid transport, cholesterol homeostasis, and lipid droplet formation, and they contribute to various signaling pathways, resulting in changes in blood-brain barrier function and increased inflammation in the brain (Badaut et al, 2024). This study focused on the NSC niches and reveals a consistent molecular mechanism of Cav-1 by which caveolae fine-tune the balance between NSC quiescence and oligodendrocyte differentiation in the V-SVZ by regulating PDGFRα protein stability, highlighting a niche-specific role of caveolae in the adult brain. Further research will be necessary because our study does not elucidate the precise mechanisms by which caveolae selectively interact with PDGFRα and transport it to lysosomes. In conclusion, we demonstrate that a V-SVZ-specific role of Cav-1 regulates NSC proliferation and subsequent differentiation into oligodendrocytes *in vivo.* Caveolae facilitate rapid lysosomal turnover of PDGFRα, ensuring proper regional fate determination and timing of activation for adult NSCs in the V-SVZ.

## Methods

### Animals

Cav-1–KO mice (C57BL/6 background) were obtained from NIBIOHN and maintained under specific pathogen-free conditions. The Cav-1–KO allele was generated by inserting a single base into exon 3 using CRISPR-Cas9, resulting in a frameshift and a premature stop codon used for genotyping. For genotyping, DNA extracted from mouse tails was amplified via PCR using the following primers: Cav-1–S2186 (forward): 5′-ATCTACAAGCCCAACAACAAGGCCA-3′, and Cav-1–R2325C (reverse): 5′-AGCAACATGCTTACCTTGACCAGGT-3′. The PCR products were then digested with AvaII (New England Biolabs, MA, USA). The wild-type allele produced an undigested 140 bp band, while the mutant allele produced a cleaved 115 bp band.Embryos from Cav-1 Het and homozygous (KO) mice at E15.5 were used to prepare primary NSC cultures (Kobayashi et al, 2019). Brains from Cav-1–Het and Cav-1–KO littermates were used for *in vivo* analysis at postnatal day 14 (P14), 2MO, and 6MO. Mice were housed in our animal facility under a 12:12 h light-dark cycle. Animal care and experiments conformed to the guidelines of the Animal Experiment Committee of Kyoto University and the University of Tokyo.

### NSC culture, quiescence induction, and *in vitro* differentiation

NSCs were cultured in proliferation medium [20 ng/mL EGF (PeproTech Inc., NJ, USA), 20 ng/mL bFGF (PeproTech Inc.), penicillin/streptomycin (P/S; Nacalai Tesque, Inc., Japan), and N-2 MAX media supplement (N-2 MAX; R&D Systems, Inc., MN, USA) in Dulbecco’s modified Eagle medium (DMEM)/F-12 (Gibco, MA, USA)] with laminin (2 µg/mL; Sigma-Aldrich, MO, USA). To induce quiescence, NSCs were seeded at 35,000–65,000 cells/cm^2^, and the proliferation medium was then replaced with quiescence medium [50 ng/mL BMP4 (Qkine Ltd., Cambridge, UK), 20 ng/mL bFGF, P/S, and N-2 MAX in DMEM/F-12] after washing three times with phosphate-buffered saline (PBS) (Martynoga et al, 2013). For RNA-seq, NSCs were cultured in quiescent medium for 4 days. *In vitro* differentiation of NSCs was induced as previously described (Imayoshi *et al*, 2013). For neuronal differentiation, NSCs were cultured in neural differentiation medium [B-27^TM^ Supplement minus vitamin A (Gibco), retinoic acid (1 μM; Nacalai Tesque, Inc.), and GlutaMAX in Neurobasal medium (Gibco)] for 3 days. For astrocyte differentiation, NSCs were cultured in astrocyte differentiation medium [N-2 MAX, LIF (20 ng/mL; Millipore, MA, USA), BMP4 (20 ng/mL), P/S, and GlutaMAX (Gibco) in DMEM/F-12] for 4 days. For OPC differentiation, NSCs were cultured in oligodendrocyte differentiation medium [N2-MAX, 10 ng/mL PDGF (R&D Systems, MN, USA), 30 ng/mL 3,3,5-triiodothyronine (T3; Sigma-Aldrich) and P/S in DMEM/F12] for 2 days. For the protein stability assay, NSCs were cultured in proliferation medium with 10 µg/mL cycloheximide (Sigma-Aldrich) for 0, 1, 2, and 4 h and then supplemented with 20 nM bafilomycin A1 (Sigma-Aldrich) or 100 nM bortezomib (Sigma-Aldrich) for 4 h as inhibitor-treated samples.

### Immunoblotting, immunocytochemistry (ICC), and immunohistochemistry (IHC)

For western blotting, the cells were washed with cold PBS and lysed on ice for 30 minutes in lysis buffer [50 mM Tris-HCl (pH 8.0), 100 mM NaCl, 5 mM MgCl₂, 0.5% Nonidet P-40, Complete™ protease inhibitor cocktail (Roche, Basel, Switzerland), 1 mM phenylmethylsulfonyl fluoride, 250 U/mL Benzonase (Sigma-Aldrich), 10 mM β-glycerophosphate, 1 mM sodium orthovanadate, 1 mM NaF, and 1 mM sodium pyrophosphate] (Kobayashi et al, 2019). After protein quantification using the DC™ Protein Assay Kit (Bio-Rad, CA, USA), the lysates were mixed with lithium dodecyl sulfate sample buffer, separated using NuPAGE™ Bis-Tris gels (Thermo Fisher Scientific, MA, USA), and transferred onto polyvinylidene fluoride membranes (Immobilon-P; Millipore) for immunoblotting. Proteins were detected using ECL™ Prime (Cytiva, MA, USA) and a FUSION Imaging System (Vilber Bio Imaging, France) (Ding et al, 2025).

For ICC, the cells were fixed with 4% paraformaldehyde (PFA) in PBS and then immunostained with primary and secondary antibodies in 1% normal donkey serum (NDS) and 0.1% Triton X-100 in PBS (PBST) after blocking in 5% NDS/PBST. Nuclei were counterstained with 4′,6-diamidino-2-phenylindole (DAPI; Sigma-Aldrich).

For IHC, mice were perfused transcardially with 4% PFA/PBS under anesthesia. Brains were post-fixed in 4% PFA/PBS and then cryoprotected in 10%, 20%, and 30% sucrose/PBS solutions. Coronal cryosections (20 μm thick) were prepared using a cryostat (CryoStar NX50, Thermo Scientific), mounted on glass slides, and stored at -80°C until use. For immunostaining, cryosections were incubated in Histo-VT One (Nacalai Tesque, Inc.) at 70°C for 20 minutes for antigen retrieval, permeabilized in 5% NDS/PBST, and incubated with primary and secondary antibodies. Nuclei were stained with DAPI.

### Antibodies

The following primary antibodies were used: anti-Cav-1 (rabbit, D46G3; Cell Signaling Technology, MA, USA; for IHC and ICC), anti-Cav-1 (rabbit, Abcam, ab2910; for western blotting), anti-DCX (rabbit, 4604; Cell Signaling Technology), anti-GFAP (mouse, G3893; Sigma-Aldrich), anti-Ki-67 (rat, SolA15; eBioscience, CA, USA), anti-Nestin (chicken; Aves, USA), anti-NeuN (mouse, MAB377; Sigma-Aldrich), anti-Olig2 (mouse, MABN50; Millipore), anti-PDGFRα (rabbit, 3164; Cell Signaling Technology), anti-Sox2 (goat, AF2018; R&D Systems), anti-S100β (rabbit, ab52642; Abcam), and anti-Tuj1 (rabbit, ab18207; Abcam) antibodies. The following secondary antibodies were used: donkey-host antibodies conjugated with Alexa Fluor 405, 488, 568, or 647 (Thermo Fisher Scientific and Abcam) and horse radish peroxidase (Promega Co., WI, USA).

### Image analysis

ICC and IHC images were acquired using a confocal microscope (Leica Stellaris 5), and three-dimensional reconstructions from z-stack images were generated using Imaris software (Oxford Instruments). Staining with Sox2, Nestin, GFAP, DCX, and S100β antibodies was performed to define the analysis area and identify cell types. To quantify Cav-1 in NSCs and astrocytes in the DG, Cav-1 signals in the vasculature were removed by masking vessel-specific Cav-1 surfaces. For the DG, quantified values were normalized either to the maximum length of the DG region identified by DAPI staining or to the DAPI surface volume. Newly generated neurons were quantified as the total volume of the DCX surface and normalized to the DAPI surface volume of the DG. The DAPI volume of the lateral SVZ was used to calculate DCX+ cells in the SVZ. The maximum lateral length (mm) of the SVZ identified by DAPI staining was used to count Olig2+ PDGFRα+ cells (OPCs) and GFAP+S100β+ cells (astrocytes). Ectopic OPCs in the ventral side of the V-SVZ were counted and normalized to the maximal lateral length (mm) of the SVZ. Quantitative analysis was performed using IHC images from at least three to eight sections per mouse.

### Quantitative mass spectrometry (MS) analysis

Total cell lysates from NSCs were prepared using the EasyPrep Mini MS Sample Prep Kit (Thermo Fisher Scientific) according to the manufacturer’s instructions. Tandem mass tag (TMT)-based proteomics was performed by labeling with TMTpro 16plex reagent (Thermo Fisher Scientific), as previously described (Morita et al, 2025). After TMT labeling, the sample channels were combined in an equal ratio, dried using a speed-vac, and resuspended in 0.1% trifluoroacetic acid. Samples were fractionated into eight fractions using a High pH Reversed-Phase Peptide Fractionation Kit (Thermo Fisher Scientific) according to the manufacturer’s protocol. Peptide (1.25 µg) from each fraction was analyzed using an EASY-nLC 1200 connected inline to an Orbitrap Fusion Lumos equipped with a FAIMS-Pro ion mobility interface (Thermo Fisher Scientific). Peptides were separated on an analytical column (C18, 1.6 µm particle size × 75 µm diameter × 250 mm; Ion Opticks AUR2-25075C18A) heated at 55°C through a column oven (Sonation) using 4 h gradients (0% to 28% acetonitrile over 240 min) at a constant flow of 300 nL/min. Peptide ionization was performed using a Nanospray Flex Ion Source (Thermo Fisher Scientific). FAIMS-Pro was set to three phases (-40, -60, and -80 CV), and the Orbitrap Fusion Lumos was operated in the data-dependent acquisition mode. Full MS scans were acquired at 120,000 resolution with a mass range of 375–1500 m/z, and the most intense ions in every 1 s were selected for MS/MS fragmentation with an HCD (higher energy collisional dissociation) of 34, isolation window at 0.7 m/z, a resolution of 50,000, and maximum injection time at 105 ms in the centroid mode The raw files were analyzed using a Sequest HT search program in Proteome Discoverer 2.4 (Thermo Fisher Scientific). MS/MS spectra were searched against the SwissProt-reviewed *Mus musculus reference* proteome (UniProt). The mass tolerances for the precursor and fragment ions were 10 ppm and 0.02 Da, respectively. Up to two missed trypsin cleavage sites were allowed. Oxidation (Met) and deamidation (Gln and Asn) were selected as variable modifications, and carbamidomethylation (Cys) and TMTpro (Lys and any N-terminus) were selected as static modifications. Peptide identification was filtered at false discovery rate < 0.01. TMT-based protein quantification was performed using the Reporter Ion Quantifier node in Proteome Discoverer 2.4. Statistical analysis and visualization were performed in R (v4.1.2) (https://www.R-project.org/) using fold-change (FC) and p-values calculated in Proteome Discoverer. Volcano plots and heatmaps were created using ggplot2 with ggrepel and complexheatmap packages in R. GO enrichment analysis was performed using the DAVID gene functional classification tool (https://davidbioinformatics.nih.gov/home.jsp).

### RNA-seq analysis

Total RNA was extracted from NSCs using the NucleoSpin RNA Kit (Takara Bio, Shiga, Japan). RNA libraries were constructed using the FLASH-seq method (Hahaut et al, 2022) and sequenced using an Illumina NextSeq 2000 platform (Illumina, CA, USA). Sequencing reads were aligned to the UCSC mm10 genome provided by Illumina iGenome using the STAR software (Dobin et al, 2013). Gene-level read counts were generated using FeatureCounts (Liao et al, 2014) and normalized to transcripts per million (TPM). Subsequent analyses were performed using RNAseqChef (Etoh & Nakao, 2023). Differential expression was assessed using DESeq2 (Wald test with Benjamini–Hochberg correction).

### Statistical analysis

Data was analyzed using GraphPad Prism. Differences between two groups were analyzed using unpaired two-tailed t-test, and differences between multiple groups were analyzed using ANOVA followed by Tukey’s test. For protein degradation kinetics, the significance between two groups were analyzed using paired t-test. Statistical significance was set at p values < 0.05 (*p < 0.05, **p < 0.01, ***p < 0.001, and ****p < 0.0001). Data in bar charts are presented as the mean, and all error bars represent the SEM.

## Acknowledgements

We thank H. Sugishita, A. Watanabe (One-stop Sharing Facility Center for Future Drug Discoveries, the University of Tokyo) for RNA-sequencing analysis; D.C. Lie, Y. Kosodo, M. Matsuda, members of the Saeki laboratory and the Matsuda laboratory for technical help and discussion: W. P. BURRIS and Editage for English text editing. This work was supported by the JST SPRING, Grant Number JPMJSP2108 (H.Z and Y. K), a Grant-in-Aid for Transformative Research Areas (Japan Society for the Promotion of Science) (23H04919 to T.K.) and by the Japan Agency for Medical Research and Development (JP20gm6410006 to T.K.).

## Data availability

Raw RNA sequencing data have been deposited to the DDBJ (Fukuda et al, 2021) with accession numbers PRJDB39769 (aNSCs and qNSCs) and PRJDB39837 (Cav-1–KO and Cav-1–HET NSCs). The mass spectrometry proteomics data have been deposited to the ProteomeXchange Consortium via the PRIDE partner repository with the dataset identifier PXD071314 (Username, Password: r9wgc8GVNXwH).

## Author contributions

**He Zhang**: Conceptualization; Resources; Formal analysis; Validation; Investigation; Visualization; Methodology; Writing—original draft. **Yusuke Kihara:** Resources; Software; Formal analysis; Validation; Investigation; Visualization; Writing—original draft. **Mitsuho Sasaki:** Resources. **Takuya Tomita:** Software; Formal analysis; Validation; Methodology; Writing—original draft. **Yasushi Saeki:** Software; Methodology; Supervision. **Taeko Kobayashi:** Conceptualization; Supervision; Formal analysis; Funding acquisition; Validation; Investigation; Visualization; Writing—original draft; Writing—review and editing.

## Disclosure and competing interests

The authors declare no competing or financial interests.

## Expanded View Figure Legends

**Figure EV1.**
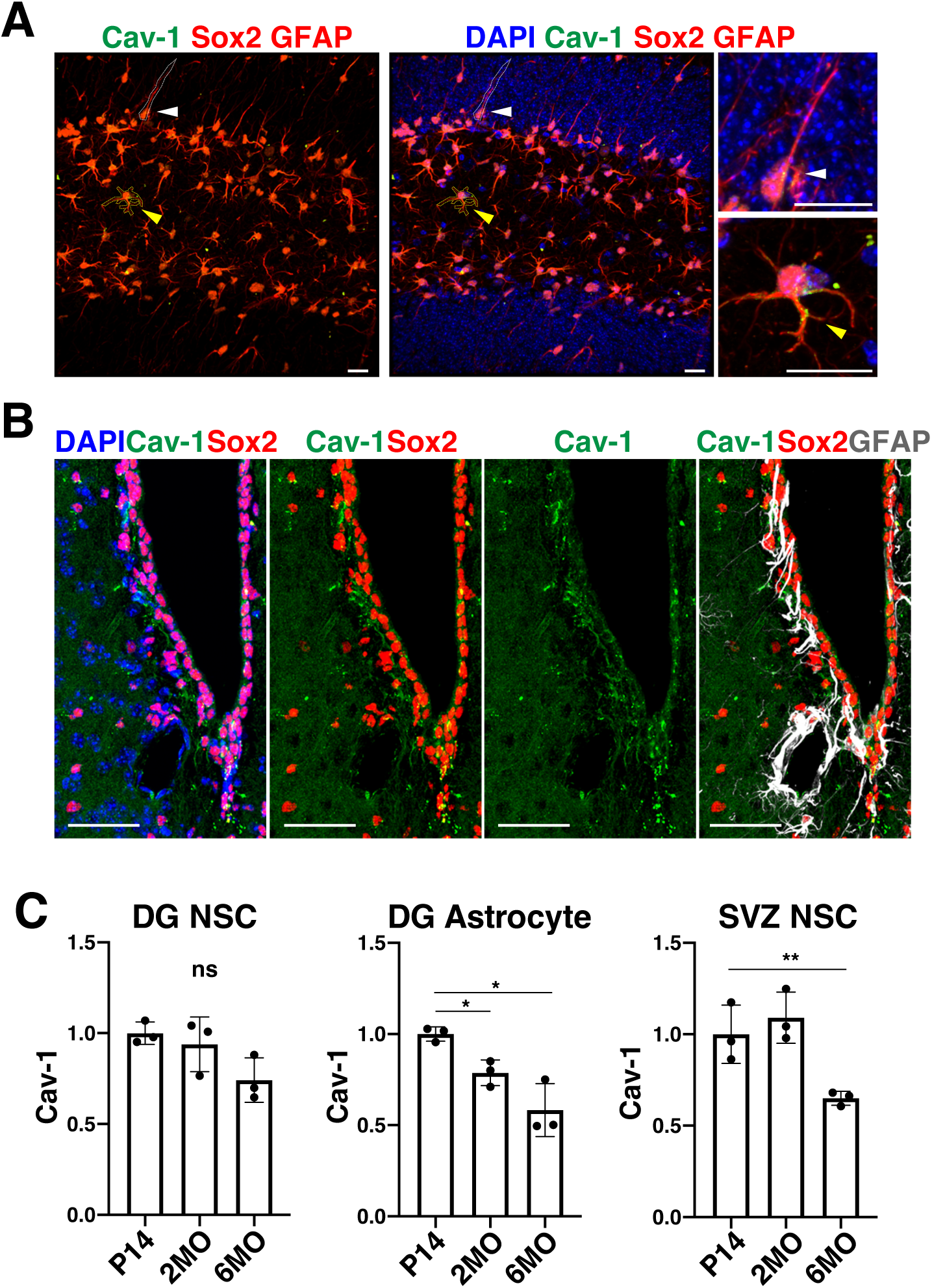
Age-dependent decline of Cav-1 expression within the DG and SVZ niches. (A) Representative confocal images of the DG showing Cav-1 expression (green) in Sox2⁺ GFAP⁺ double-positive cells (red), including NSCs (white dashed line, white arrowhead) and astrocytes (yellow dashed line, yellow arrowhead) at 2MO. Nuclei are labeled with DAPI (blue). Enlarged views of NSC (upper, white arrowhead) and astrocyte (lower, yellow arrowhead) are in the right panels. Scale bars, 20 µm. (B) Representative images of the SVZ showing Cav-1 (green), Sox2 (red), GFAP (white), and DAPI (blue) at 2MO. Scale bars, 50 µm. (C) Quantification of Cav-1 intensity in DG NSCs (left), DG astrocytes (middle), and SVZ NSCs (right) at P14, 2MO, and 6MO. Values are normalized to the mean intensity at P14. Data represent SD from three sections in one mouse per age group. One-way ANOVA with Tukey’s test (*p < 0.05; **p < 0.01; and ns, not significant).

**Figure EV2.**
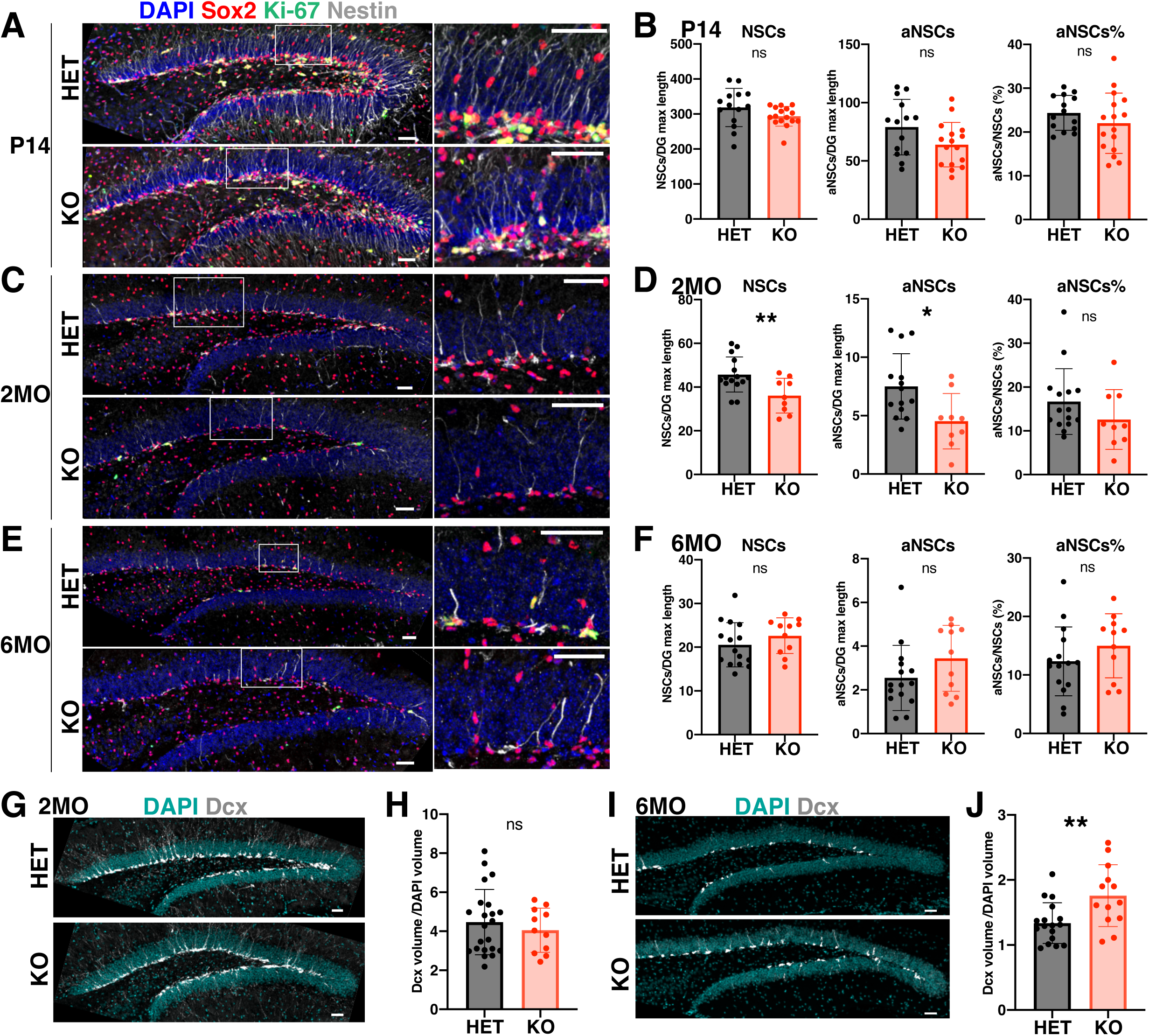
Cav-1 deletion does not significantly impact NSC maintenance in the DG across ages from P14 to 6MO. (A, C, E) Representative confocal images of the DG from Cav-1–HET and Cav-1–KO mice at P14 (A), 2MO (C), and 6MO (E), stained for Sox2⁺ (red), Ki-67⁺ (green), and Nestin⁺ (white) and with DAPI (blue). Right panels show high-magnification views of the boxed regions in the left panels. Scale bars, 100 µm (left) and 50 µm (right). (B, D, F) Quantification of DG NSCs (Sox2⁺Nestin⁺/DG maximal length), active NSCs (aNSCs; Sox2⁺Nestin⁺Ki-67⁺/DG maximal length), and the percentage of aNSCs within total NSCs (aNSCs/NSCs) at P14 (B), 2MO (D), and 6MO (F). Mean ± SD; n = 3 mice per group; unpaired two-tailed t-test (*p < 0.05; **p < 0.01; and ns, not significant). (G-J) Representative DCX⁺ staining of newly generated neurons in the DG at 2MO (G) and 6MO (I), with quantification of DCX⁺ volume normalized to DAPI⁺ volume of the DG at 2MO (H) and 6MO (J). Scale bars, 100 µm. Mean ± SD; n = 3 mice per group; unpaired two-tailed t-test (**p < 0.01).

**Figure EV3.**
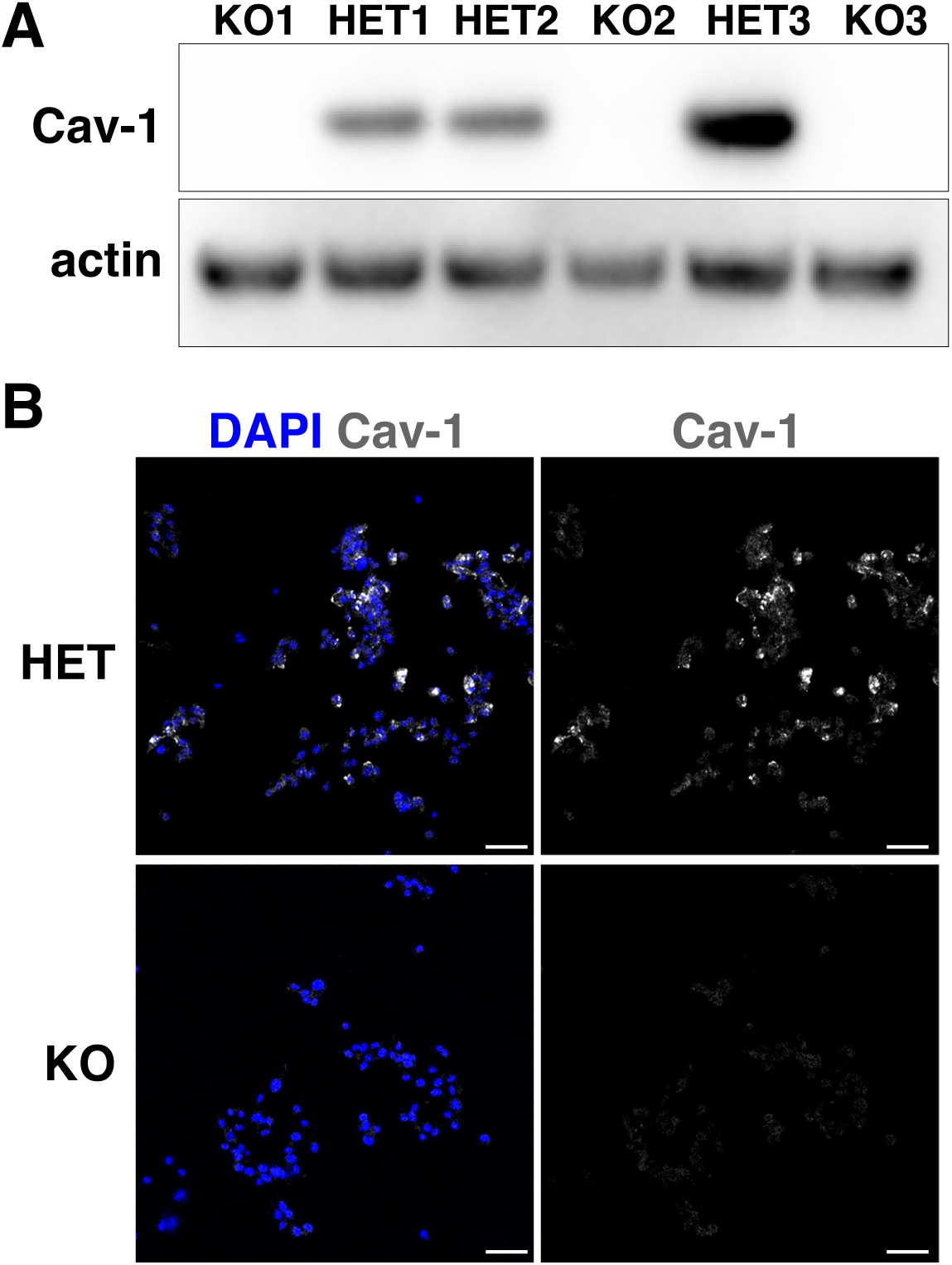
Validation of Cav-1–KO in NSCs. (A) Western blot analysis of Cav-1 protein expression. Actin was used as a loading control. KO1/KO2/KO3 and HET1/HET2/HET3 represent independent NSCs from different embryos. (B) Immunocytochemistry of Cav-1 in HET and KO NSCs. Nuclei are labeled with DAPI (blue). Scale bars, 50 µm.

**Figure EV4.**
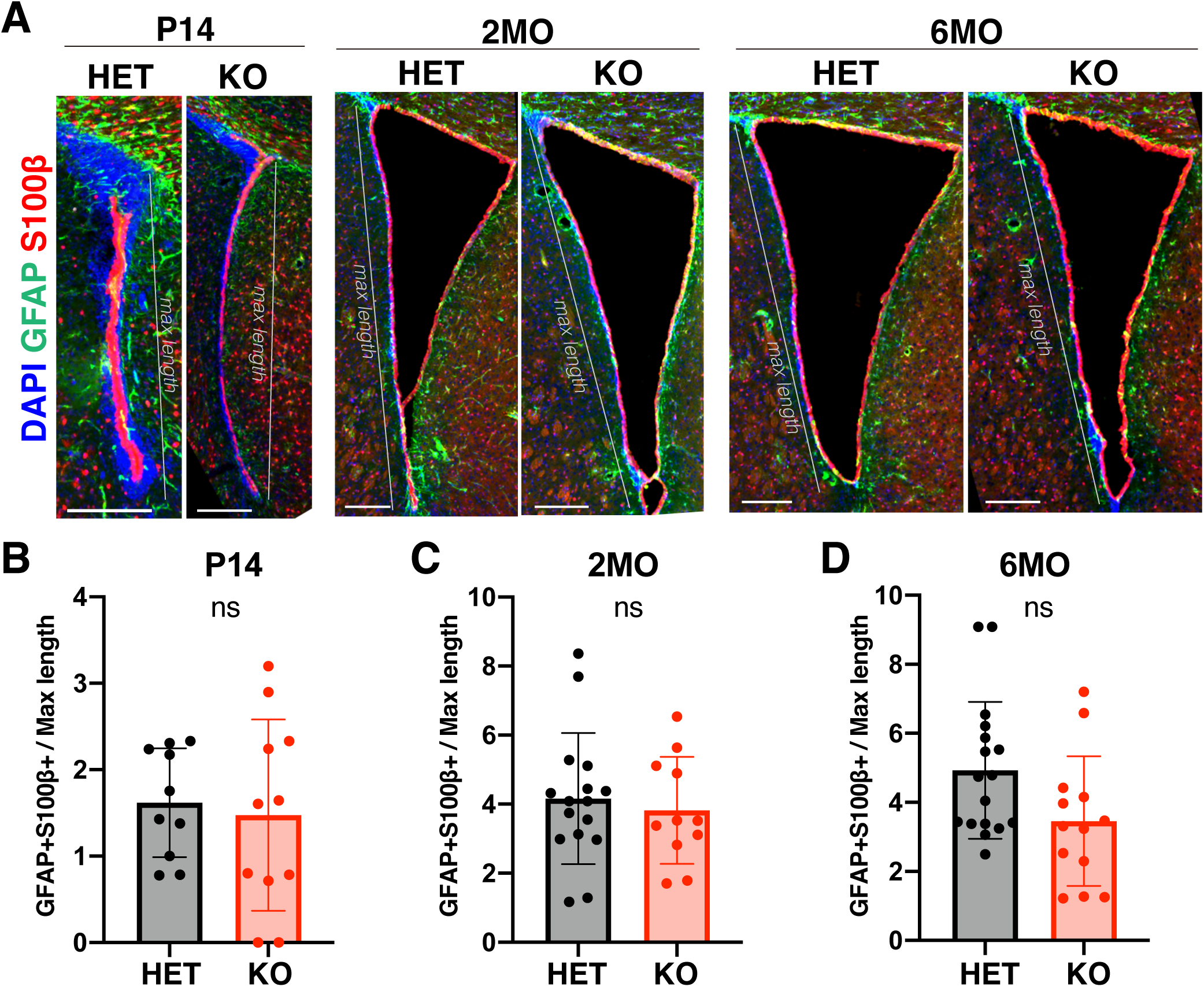
Cav-1 deletion does not significantly impact astrocyte differentiation in the V-SVG across ages from P14 to 6MO. (A) Representative confocal images of the V-SVZ from Cav-1–HET and Cav-1–KO mice at P14, 2MO, and 6MO, stained for GFAP⁺ (green), S100β⁺ (red), and with DAPI (blue). Grey lines indicate the maximal length of the lateral V-SVZ used for quantification. Scale bars, 200 µm. (B, C, D) Quantification of astrocytes (GFAP⁺S100β⁺/SVZ maximal length) in the lateral V-SVZ at P14 (B), 2MO (C), and 6MO (D). Astrocyte numbers were counted within 100 µm from the lateral side of the ventricle. Mean ± SD; n = 3 mice per group; unpaired two-tailed t-test (ns, not significant).

**Figure EV5.**
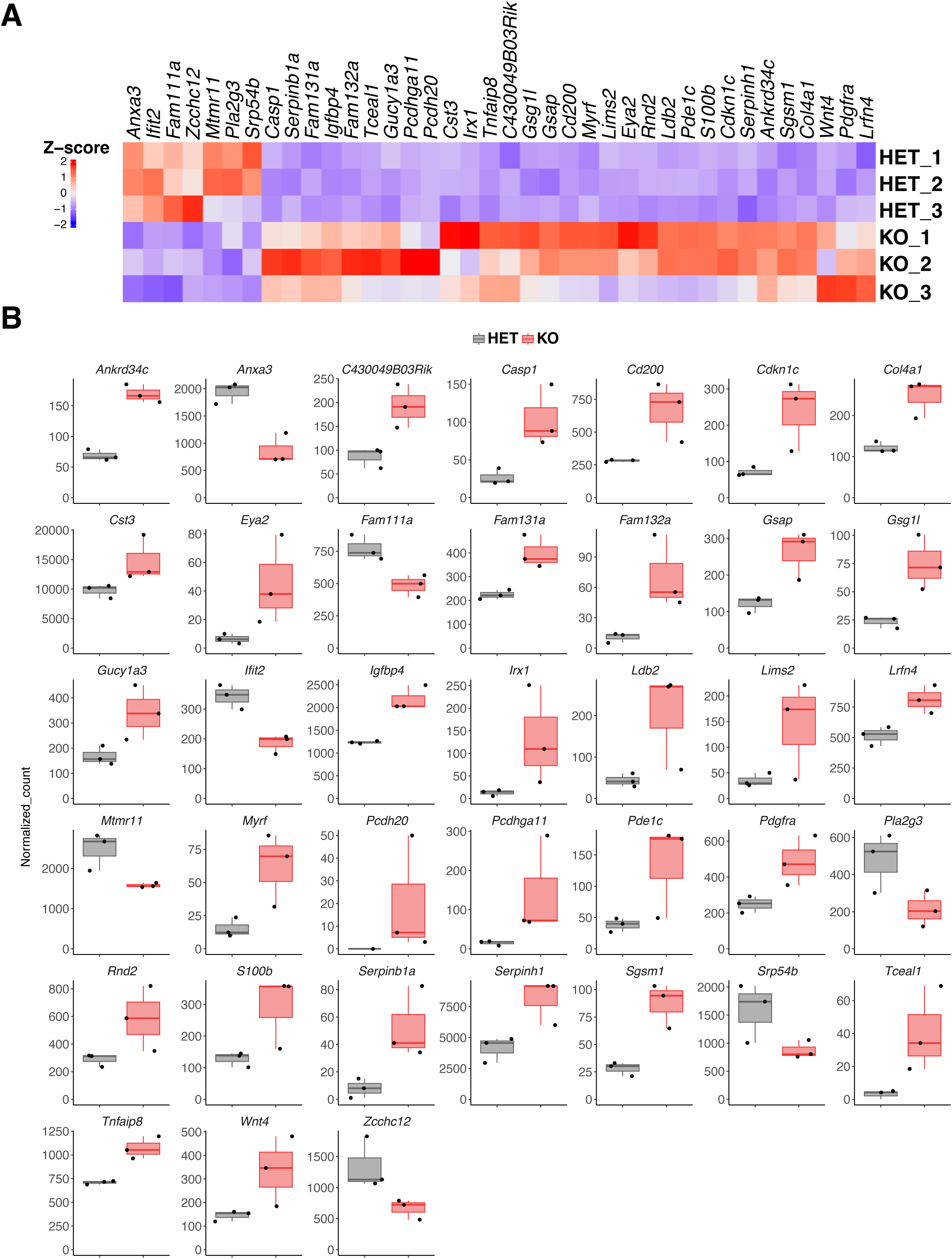
Transcriptome profiles in Cav-1–HET and Cav-1–KO NSCs. (A) Heatmap showing z-score–normalized expression of up- and down-regulated genes in Cav-1–KO NSCs compared with control HET NSCs (HET_1–3, KO_1–3; n = 3). Genes were selected from the Cav-1–regulated transcript set identified via RNA-seq (Fig. 5A). DESeq2, Padj < 0.05, Fold change ≥ |1.5|. (B) Boxplots showing normalized RNA-seq counts for each gene in Cav-1–HET (grey) and Cav-1–KO (red) NSCs. Boxes show interquartile range with median (n = 3). All genes shown are significantly different between Cav-1–HET and Cav-1–KO NSCs according to DESeq2 (Padj < 0.05).

## Expanded View Tables

**Table EV1 – Proteome data comparing aNSCs and qNSCs.**

(A) Protein abundances for aNSCs and qNSCs identified by quantitative TMT-MS, used to create the heatmap in Fig. 1A (n=2).

(B) Fold change (FC) of protein abundances in qNSCs relative to those in aNSCs, and the p-value calculated by Proteome Discoverer, used to generate the volcano plot in Fig. 1B. Some factors, such as Ccdc3, Cep162, Disp3, Kcna4, and Mutyh, were excluded in this volcano plot table because p-values could not be calculated.

**Table EV2 – Gene ontology results for factors at least twofold enriched in qNSCs compared to aNSCs in Table EV2.**

**Table EV3 – Transcriptome results comparing aNSCs (A1-3) and qNSCs (Q1-3).**

**Table EV4 – Transcriptome results in Cav-1–HET (HET1-3) and Cav-1–KO (KO1-3) NSCs.**

## Notes

### Competing Interest Statement

The authors have declared no competing interest.

